# Neurotransmitters Contribute Structure-Function Coupling: Evidence from Grey Matter Volume (GMV) and Brain Entropy (BEN)

**DOI:** 10.1101/2024.09.07.611832

**Authors:** Donghui Song, Ze Wang

**Author notes:** Corresponding author: Donghui Song, No.19, Xinjiekouwai St, Haidian District, Beijing, 100875, CN. Corresponding author: Ze Wang, Ph.D., 670 W Baltimore St, Baltimore, MD, 21201, USA.

## Abstract

The emergence of human brain function from macroscopic anatomical structures and its relationship with the coupling of structure and function have long been pivotal questions in neuroscience. Neurotransmitter receptors are critical in signal transmission, regulating brain function, and potentially enhancing the coupling between brain structure and function. Recent research suggests that neurotransmitter systems facilitate this coupling between structural and functional networks. However, the mechanisms of how neurotransmitters adapt to local anatomical structures and drive functional emergence are not yet fully understood. This study explores these mechanisms using gray matter volume (GMV) and brain entropy (BEN). BEN reflects the irregularity, unpredictability, and complexity of brain activity, and functional MRI (fMRI)-based BEN has identified its distribution in the normal human brain. BEN is correlated to cognitive and task performance, and aberrant BEN patterns link to various brain disorders. Notably, BEN can indicate neuroplasticity through non-invasive brain stimulation, and pharmacological intervention. We analyzed structural imaging data, as well as 7T resting-state and movie-watching fMRI data from 176 participants in the Human Connectome Project (HCP), to calculate GMV, resting-state BEN (rsBEN), and movie-watching BEN (mvBEN). By integrating data from the publicly available neurotransmitter receptors database, we evaluated the role of neurotransmitters in the coupling between GMV and rsBEN, GMV and mvBEN, and rsBEN and mvBEN. Our findings reveal that neurotransmitters significantly contribute to structural and functional coupling and influence the transition from task-free to movie-watching conditions.

## 1. Introduction

Neurotransmitter receptors modulate cell firing rates by regulating excitability or inhibition, thus driving the propagation of electrical signals between synapses, facilitating synaptic plasticity, and regulating neural states to achieve various functions. These receptors exhibit a wide range of structures and functions. They can be categorized as either ionotropic or metabotropic, may be made up of multiple subunits, and can influence neural circuits in either a facilitative or inhibitory manner. Furthermore, they are associated with different downstream biochemical pathways. Recently Hansen et al (2022) utilized PET imaging data from 1200 individuals to construct maps of nine major neurotransmitter systems and determined their relationships with structural and functional networks (Hansen, Shafiei et al. 2022). Their findings showed that neurotransmitter distribution contributes to both structural and functional networks and promotes their coupling. However, how spatial distributions of neurotransmitter receptors relate to brain structure and contribute to brain function at the local brain activity level remains unknown. In the study, we attempt to address the contribution of neurotransmitters to regional structure-function coupling, and then increase our understanding of the link between structure and function with the chemoarchitecture of the brain. We investigate the coupling between gray matter volume (GMV) and resting-state brain entropy (rsBEN), and movie-watching brain entropy (mvBEN) to provide specific answers to this question. We selected these metrics based on the following considerations: GMV is the most used structural brain measurement metric, which has been employed in brain development and brain disorders (Ashburner and Friston 2000, Good, Johnsrude et al. 2001, Mechelli, Price et al. 2005). BEN measures the irregularity, disorder, and complexity of brain activity (Wang, Li et al. 2014). The rsBEN is associated with neurocognitive (Wang 2021, Del Mauro and Wang 2024), task performance (Lin, Chang et al. 2022), GMV (Del Mauro and Wang 2024), neuromodulation (Song, Chang et al. 2019, Liu, Song et al. 2024, Song, Deng et al. 2024), hormones (Song and Wang 2024) and various brain disorders (Chang, Song et al. 2018, Xue, Yu et al. 2019, Liu, Song et al. 2020, Jiang, Cai et al. 2023, Del Mauro, Sevel et al. 2024, Dong-Hui Song 2024). Our recent studies have also identified BEN associated with movie-watching (Song and Wang 2024). Importantly, there is a progressive relationship between these functional measurements, from task-free to movie-watching that across various cognitive domains. This allows for the consideration of the relationship between neurotransmitters and functional brain activation at different levels.

## 2. Methods

All participants from the Human Connectome Project (HCP) 7T release. One hundred seventy-six (106 female) out of 184 participants completed T1w, four resting state (REST) runs, and four movie-watching (MOVIE) runs over two days. All participants were healthy individuals between 22 and 36 years (mean age = 29.4, standard deviation = 3.3). The structural images were scanned on a 3T Siemens Skyra scanner using a standard 32-channel head coil at Washington University. The functional images were acquired on a 7 Tesla Siemens Magnetom scanner at the Center for Magnetic Resonance Research at the University of Minnesota. Detailed information about data acquisition and preprocessing can be found in (Glasser, Sotiropoulos et al. 2013, Van Essen, Smith et al. 2013), also see (Song and Wang 2024). The structural images were preprocessed including distortion correction and bias field correction by HCP named “T1w_acpc_dc_restore_brain.nii.gz”. GMV was computed by the FSL pipeline (Smith, Jenkinson et al. 2004, Douaud, Smith et al. 2007) following FSLVBM protocol (https://fsl.fmrib.ox.ac.uk/fsl/fslwiki/FSLVBM), fslvbm_2_template was used to create the study-specific GM template, fslvbm_3_proc was used to register all GM images to the study-specific template non-linearly. The detailed guide of FSLVBM is available at https://fsl.fmrib.ox.ac.uk/fsl/fslwiki/FSLVBM/UserGuide.

BEN maps were calculated from our previous study (Song and Wang 2024). The GMV maps and BEN maps were parcellated to 400 parcels according to atlas from (Schaefer, Kong et al. 2018) by neuromaps (Markello, Hansen et al. 2022). Neurotransmitter receptor and transporter maps from https://github.com/netneurolab/hansen_receptors/tree/main/data/PET_nifti_images, and also were parcellated to 400 parcels.

## 3. Results

### 3.1 Spatial correlation of neurotransmitter receptors with GMV, rsBEN and mvBEN

We first evaluated the spatial correlation between 19 neurotransmitter receptor or transporter maps with GMV, rsBEN, and mvBEN. For each neurotransmitter receptor or transporter map, we conducted Pearson’s correlation analyses with GMV, rsBEN, and mvBEN from each participant. Similarly, we also assessed the spatial correlations among GMV, rsBEN, and mvBEN.

The results revealed significant spatial correlations between GMV, rsBEN, and mvBEN. The average spatial correlation coefficients from 176 participants were as follows: r (400) =0.248±0.121(p<0.001, n=176) between GMV and rsBEN, *r* (400) =0.245±0.109 (p<0.001, n=176) between GMV and mvBEN, and *r* (400) =0.863±0.075(p<0.001, n=176) between rsBEN and mvBEN. The spatial correlations between GMV, rsBEN, and mvBEN and various neurotransmitter receptor or transporter maps differed. Specifically, GMV showed significant positive correlations with 5HT1a (*r* (400)= 0.379±0.069, *p*<0.001, n=176), 5HTT(*r* (400)= 0.444±0.057, *p*<0.001, n=176), D1(*r* (400)= 0.473±0.068, *p*<0.001, n=176),D2(*r* (400)= 0.233±0.060, *p*<0.001, n=176), DAT(*r* (400)= 0.562±0.072, *p*<0.001, n=176), H3(*r* (400)= 0.278±0.056, *p*<0.001, n=176), MU(*r* (400)= 0.312±0.099, *p*<0.001, n=176), NMDA(*r* (400)= 0.491±0.062, *p*<0.001, n=176), and VAChT(*r* (400)= 0.442±0.040, *p*<0.001, n=176), while it had a significant negative correlation with 5HT1b(*r* (400)=-0.337±0.060, *p*<0.001, n=176). rsBEN exhibited significant positive correlations with 5HT1a(*r* (400)= 0.279±0.092, *p*<0.001, n=176), 5HT2a (*r* (400)= 0.262±0.084, *p*<0.001, n=176), 5HT4(*r* (400)= 0.196±0.084, *p*<0.001, n=176), 5HT6(*r* (400)= 0.249±0.102, *p*<0.001, n=176), D1(*r* (400)= 0.318±0.108, *p*<0.001, n=176), D2(*r* (400)= 0.185±0.092, *p*<0.001, n=176), DAT(*r* (400)= 0.31±0.133, *p*<0.001, n=176), GABAa(*r* (400)= 0.192±0.086, *p*<0.001, n=176), MU(*r* (400)= 0.201±0.105, *p*<0.001, n=176), and NMDA(*r* (400)= 0.332±0.117, *p*<0.001, n=176). mvBEN was significantly positively correlated with 5HT1a(*r* (400) = 0.341±0.093, *p*<0.001, n=176), 5HTT(*r* (400) = 0.492±0.044, *p*<0.001, n=176), D1(*r* (400) = 0.452±0.086, *p*<0.001, n=176), D2(*r* (400) =0.252±0.062, *p*<0.001, n=176), DAT(*r* (400) = 0.602±0.060, *p*<0.001, n=176), H3(*r* (400) =0.188±0.074, *p*<0.001, n=176), NMDA(*r* (400)= 0.526±0.050, *p*<0.001, n=176), and VAChT(*r* (400) = 0.437±0.037, *p*<0.001, n=176), but showed a significant negative correlation with 5HT1b (*r* (400)= -0.342±0.062, *p*<0.001, n=176) (see Fig1).

**Fig1.**
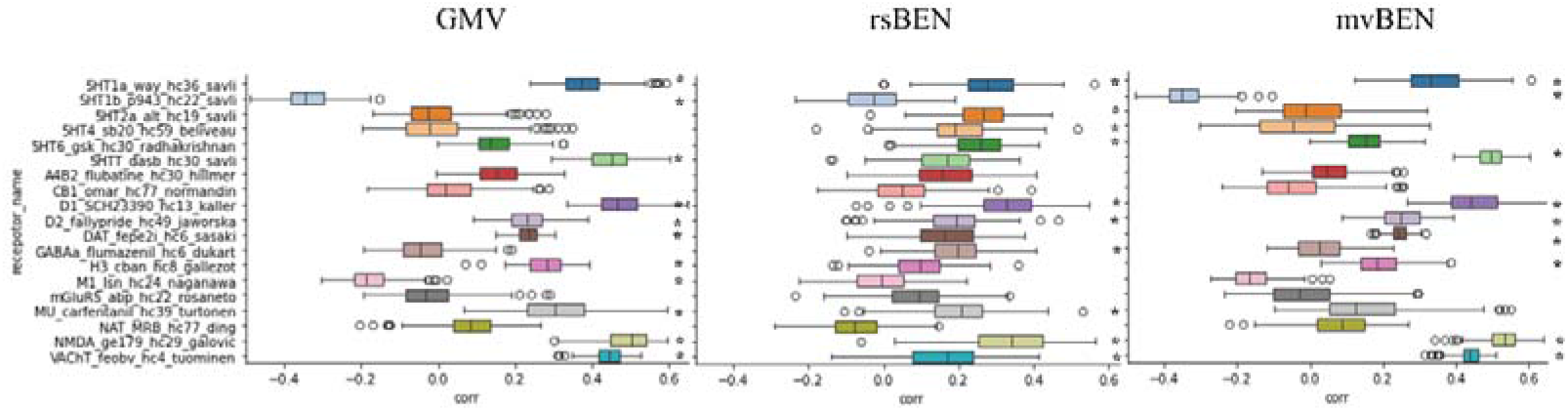
Spatial correlation of neurotransmitter receptor or transporter maps with GMV, rsBEN and mvBEN. The y-axis represents the names of the receptors or transporter map, while the x-axis displays the correlation coefficients. From left to right, the plot shows the spatial correlations between the receptors or transporter maps and GMV, rsBEN, and mvBEN, respectively. * indicates significance level p<0.001.

### 3.2 Receptor distributions contribute structure–function coupling

To assess the coupling between receptor distribution and structural and functional features, we constructed receptor similarity maps based on Hansen et al (Hansen, Shafiei et al. 2022). Similarly, we created distribution similarity maps for GMV, rsBEN, and mvBEN. For the receptor similarity map, please refer to (Hansen, Shafiei et al. 2022). The similarity maps for GMV, rsBEN, and mvBEN are shown in Fig 2A. We calculated the correlations between these similarity maps using the lower triangular values. The distribution similarity correlation coefficients are as follows: r (79800) =0.248 (p<0.001) between GMV and rsBEN, r(79800)=0.266 (p<0.001) between GMV and mvBEN, r(79800)=0.733(p<0.001).We also performed principal component analysis (PCA) to reduce the dimensionality of the similarity maps for GMV, rsBEN, and mvBEN. The highest loading component, PC1, is shown in Fig 2A. We calculated the correlations between these PC1 scores, with the following results: r (400) =0.485 (p<0.001) between GMV and rsBEN, r (400) =0.478 (p<0.001) between GMV and mvBEN, r (400) =0.912 (p<0.001) between rsBEN and mvBEN (see Fig2A).

**Fig2.**
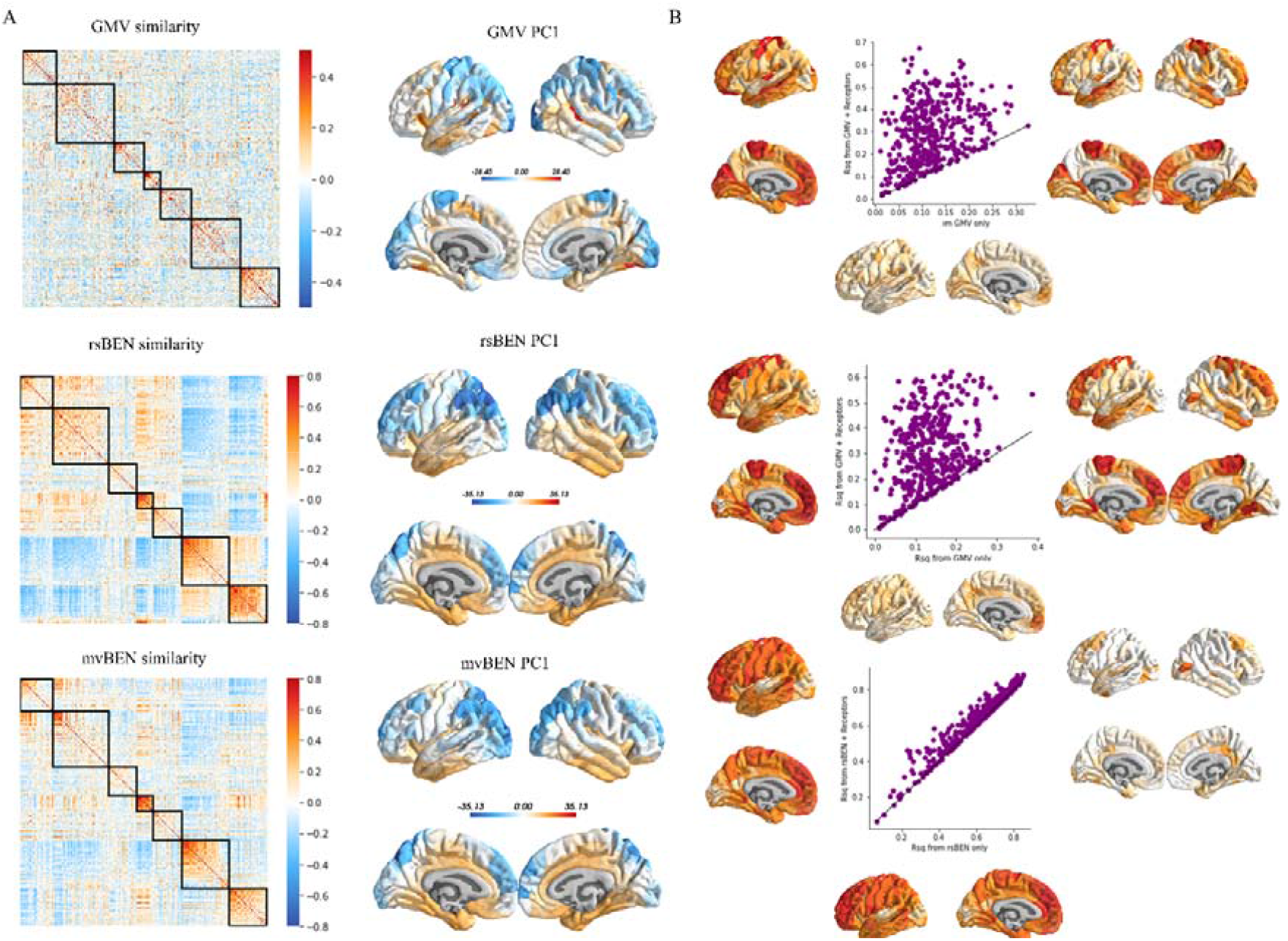
Receptor distributions contribute structure–function coupling. A. For each pair of brain regions, GMV and BEN values from each participants (n=176) are correlated (Pearson□s r) to construct the GMV and BEN similarity matrix (ordered according to the Yeo–Krienen intrinsic networks (Yeo, Krienen et al. 2011): frontoparietal, default mode, dorsal attention, limbic, ventral attention, somatomotor and visual. Left: From top to bottom are the similarity matrices for GMV, rsBEN, and mvBEN. Right: from top to bottom are the scores of PCA1 for the similarity matrices of GMV, rsBEN, and mvBEN. B. Regional structure–function or rest-task coupling was computed as the fit (R^2^) between GMV and rsBEN or mv BEN. From top to bottom show that regional structure–function coupling was computed as the fit (R^2^) between GMV and rsBEN, regional structure–function or rest-task coupling was computed as the fit (R^2^) between GMV and mvBEN and regional rest-task coupling was computed as the fit (R^2^) between rsBEN and mvBEN. For each group, structure–function or rest-task coupling at each brain region is plotted when receptor similarity is excluded (x-axis) and included (y-axis) in the model, the R^2^ difference between receptor similarity is excluded (x-axis) and included (y-axis) in the model.

We used the adjusted R^2^ of a simple linear regression model as structure–function coupling at every brain region, then regional receptor similarity was included as an independent variable, to assess how receptor information to change structure–function coupling. We also used the adjusted R^2^ of a simple linear regression model as rest (rsBEN)–task (mvBEN) coupling at every brain region, then regional receptor similarity was included as an independent variable, to assess how receptor information to change rest-task coupling. The paired sample t-test was used to assess whether adding receptor information significantly altered the R^2^values, which indicate the degree of structural-functional coupling. We found that including receptor profiles as an input variable alongside GMV significantly improves the prediction of regional rsBEN (t=26.89,p<0.001) and mvBEN (t<26.33,p<0.001) in visual cortex, temporal cortex, dorsal prefrontal cortex and the paracentral lobule, while including receptor profiles as an input variable alongside rsBEN significantly improves the prediction of regional mvBEN (t<15.94,p<0.001) in visual cortex, temporal cortex, posterior cingulate cortex and dorsal lateral prefrontal cortex (see Fig2B).

### 3.2 Single receptor contributes structure–function coupling

To assess the contribution of individual receptors or transporters to structural and functional coupling, we also used the adjusted R^2^of a linear model as the indicator of structural-functional coupling. In this analysis, instead of using similarity matrix, we utilized the values of GMV, rsBEN, and mvBEN for each participant. Each receptor was included as an independent variable to evaluate its effect on changes in structural-functional coupling.

The results indicate that the neurotransmitter receptors contributing to the coupling between GMV and rsBEN are like those contributing to the coupling between GMV and mvBEN and these neurotransmitter receptors include 5HT1a, 5HT1b, 5HTT, D1, MU, VAChT, NMDA, and H3. However, no single neurotransmitter receptor had a significant impact on the coupling between rsBEN and mvBEN (see Fig3).

**Fig3.**
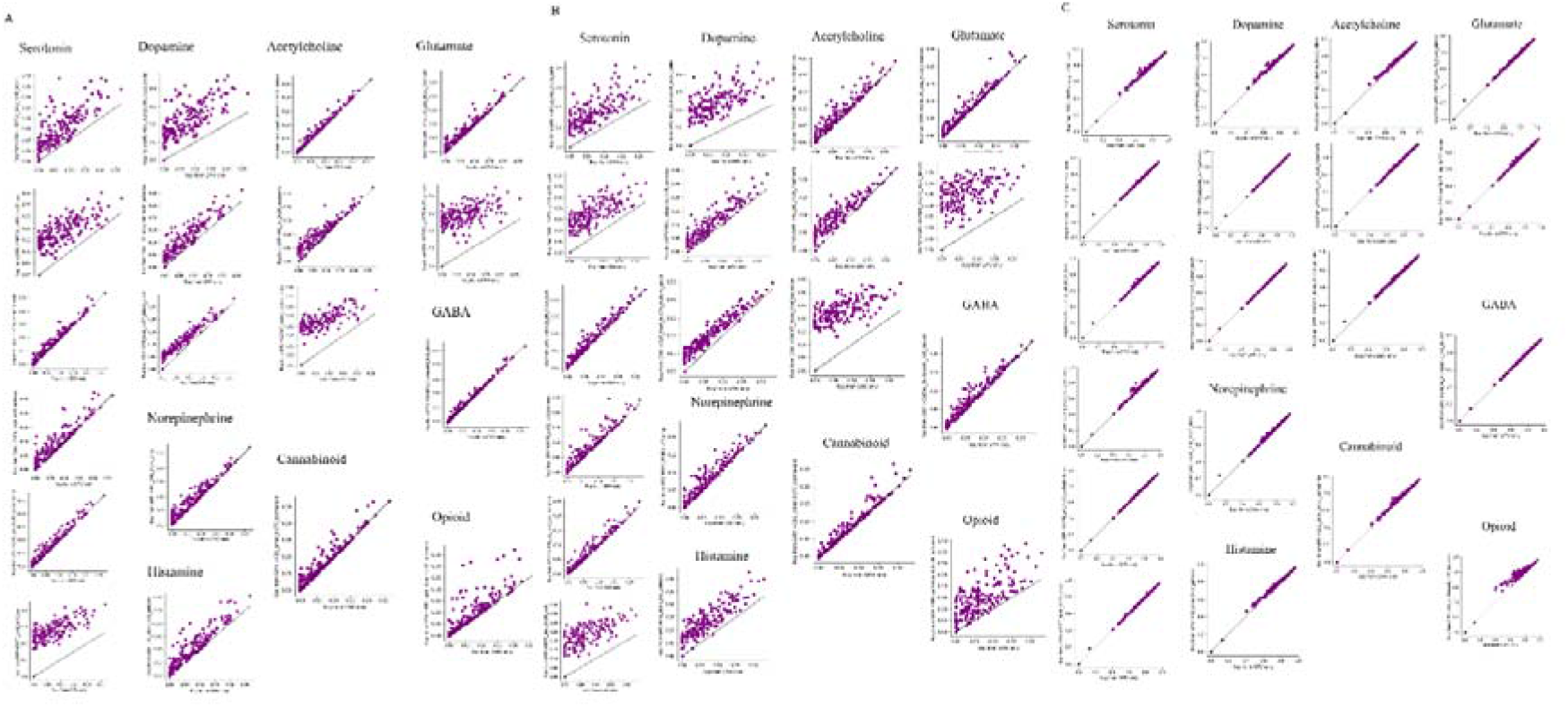
Single receptor contributes structure–function coupling. A Single receptor contribute GMV–rsBEN coupling, B Single receptor contribute GMV–mvBEN coupling, C Single receptor contribute rsBEN-mvBEN coupling.

## 4. Discussion

In this study, we assessed the spatial correlations between major neurotransmitter receptors and GMV, rsBEN, and mvBEN. We also evaluated the impact of neurotransmitter distribution on structural and functional coupling, as well as the strength of coupling for each individual neurotransmitter receptor. Our use of GMV and BEN provides new insights into the contributions of neurotransmitters to structural-functional coupling.

Recent studies have analyzed the relationship between changes in BEN before and after non-invasive interventions and neurotransmitter receptors (Liu, Song et al. 2024). However, these analyses often rely on small sample sizes and shorter scanning durations and lack fundamental spatial pattern results for neurotransmitter receptors and BEN from normal adults that limited to restricts the interpretability of the findings. In this study, we assessed the spatial correlations between each neurotransmitter receptor and GMV, rsBEN, and mvBEN. Our results provide a foundation for future applications of neurotransmitter spatial correlations with GMV, rsBEN, and mvBEN.

In the analysis of how neurotransmitters enhance structural and functional coupling, the brain regions where neurotransmitters contribute significantly to coupling overlap with those identified in previous structural and functional network analyses (Hansen, Shafiei et al. 2022). These regions include the visual cortex, temporal cortex, and paracentral lobule, as well as the dorsal prefrontal cortex. This result suggests that, whether from network-based or localized analyses, neurotransmitters contribute to structural and functional coupling in similar brain regions. In this study, we not only provide new evidence for how neurotransmitters facilitate structural and functional coupling, but we also offer potential evidence for the contribution of neurotransmitters from resting state to task state. However, since neurotransmitter distributions are static and we used averaged results from multiple movie data for mvBEN, this effect was relatively modest. In future formal studies, we will incorporate task-state data to build a more comprehensive understanding of how neurotransmitters contribute to structural-functional coupling and their contributions from task-free to specific task conditions.

**Let me (Donghui Song) continue refining this work after I complete my doctoral thesis defense!**

## Acknowledgements

MRI data were provided by the Human Connectome Project, WU-Minn Consortium (Principal Investigators: David Van Essen and Kamil Ugurbil; 1U54MH091657) funded by the 16 NIH Institutes and Centers that support the NIH Blueprint for Neuroscience R esearch; and by the McDonnell Center for Systems Neuroscience at Washington Universi ty. We thank Justine Y. Hansen et al for sharing neurotransmitter receptor maps and c odes.

## Data and code availability

All neurotransmitter receptor maps can be found at https://github.com/netneurolab/hansen_receptors/tree/main/data/PET_nifti_images.

All structural and functional MRI data are available at https://www.humanconnectome.org/.

BENtbx is available at https://www.cfn.upenn.edu/zewang/BENtbx.php.

Customed codes and further updates related to the study will be available at https://github.com/donghui1119/Brain_Entropy_Project/tree/main/Neurotransmitter_BEN (upon pu blication of the manuscript).

## CRediT authorship contribution statement

Donghui Song: conceptualization, data analysis, visualization, manuscript drafting, and editing. Ze Wang: conceptualization, manuscript editing, supervision, project admi nistration.

